# TlyA is a 23S and 16S 2′-O-methylcytidine methyltransferase important for ribosome assembly in *Bacillus subtilis*

**DOI:** 10.1101/2025.04.21.649808

**Authors:** Jennie L. Hibma, Lia M. Munson, Joshua D. Jones, Taylor M. Nye, Kristin S. Koutmou, Lyle A. Simmons

## Abstract

Ribosomal RNA (rRNA) is methylated in organisms ranging from bacteria to metazoans. Despite the pervasiveness of rRNA methylation in biology, the function of rRNA methylation on ribosome function is poorly understood. In this work, we identify a biological function for the rRNA 2′-O-methylcytidine methyltransferase TlyA, conserved between *Bacillus subtilis* and *Mycobacterium tuberculosis (Mtb)*. The *tlyA* deletion in *B. subtilis* confers a cold sensitive phenotype and resistance to aminoglycoside antibiotics that target the 16S rRNA. We show that *ΔtlyA* cells have ribosome assembly defects characterized by accumulation of the 50S subunit. Using a genetic approach and based on sequence alignments with other rRNA methyltransferases we tested the importance of potential catalytic residues and S-adenosyl-L-methionine (SAM) cofactor binding sites. We show that TlyA shares the common rRNA methyltransferase catalytic triad KDK and a SAM binding motif GxSxG which differs from *Mtb* TlyA. Together our work demonstrates that *B. subtilis tlyA* is critical for ribosome assembly and we identify key residues for TlyA function *in vivo*. Since *E. coli* lacks TlyA or a functional equivalent, our work highlights key differences in ribosome maturation between *B. subtilis, Mtb* and more divergent Gram-negative bacteria providing new insight into translation and antibiotic resistance mechanisms.

## INTRODUCTION

Ribosomes are one of the most abundant, complex, and efficient ribonucleoprotein structures found in the cell (Staley & Woolford, 2009). They function by translating messenger RNA (mRNA) transcribed from coding sequences in genomic DNA into polypeptides for use in virtually all cellular functions (Brenner et al., 1961). Mature ribosomes decode mRNA at a rate of approximately 50 nucleotides/second (nt s^-1^) (Garrett, 1999; Johnson et al., 2020). To translate proteins with such high speed and accuracy, proper ribosome architecture must be efficiently achieved and maintained.

The first step towards attaining functional ribosome architecture in bacteria is accomplished by a series of endonucleases that cleave ribosomal RNA (rRNA) transcripts which are transcribed together from multiple *rrn* operons in the bacterial chromosome (Giuliano & Engl, 2021; Herskovitz & Bechhofer, 2000). The cleaving of intergenic regions yields mature 16S, 23S, and 5S rRNAs which then fold independently or with the help of ribosome biogenesis factors to prevent misfolding and ensure assembly with the appropriate ribosomal proteins (r-proteins) (Davis & Williamson, 2017; Shajani et al., 2011). In addition to folding and binding r-proteins, rRNA maturation also includes the addition of post-transcriptional modifications. Ribosomal RNA modifying enzymes, predominantly methyltransferases (MTases) and pseudouridine synthases, are responsible for catalyzing the chemical modification of rRNA nucleobases or ribose sugars (Boccaletto et al., 2022; Decatur & Fournier, 2002). Ribosomal RNA methyltransferases (MTases) methylate either the 2′-hydroxyl group of the ribose sugar or the nitrogenous base, using S-adenosyl-L-methionine (SAM) to transfer a methyl group to their target substrate recognized by sequence or secondary structure (Ayadi et al., 2019; Decatur & Fournier, 2002). Methylation modifications contribute to rRNA folding by creating steric hinderance or directing hydrogen bonding and hence are an important factor in the formation of functional ribosomal subunits (Sharma & Entian, 2022). The secondary structure of rRNA is important for the cooperative binding of r-proteins and formation of the peptidyl transferase catalytic center of the ribosome for efficient mRNA decoding (Rodnina et al., 2020). Indeed, many modifications are clustered in the functionally important regions of the ribosome used for decoding, peptidyl transfer, tRNA binding, and the peptide exit tunnel (Antoine et al., 2021; Chow et al., 2007). Through the cumulative actions of these proteins, the final mature bacterial ribosome (70S) is formed composed of the large subunit (50S) containing 23S and 5S rRNAs and 33 r-proteins and the small (30S) subunit containing the 16S rRNA and 21 r-proteins (Aoyama et al., 2022).

Much of our knowledge on bacterial rRNA methyltransferases and their modifications come from studies in *E. coli* which contain 24 rRNA methylations incorporated by 23 different methyltransferases (Pletnev et al., 2020). Comparatively, only five rRNA methyltransferases have been studied in *B. subtilis*, the Gram-positive model organism, RsmG, RlmCD, RlmP, RlmQ, and TlyA. Only two of these, RlmP and RlmQ, have been phenotypically characterized (Desmolaize et al., 2011; Nishimura et al., 2007; Popova et al., 2024; Roovers et al., 2022; Wolff et al., 2024). Recently, the modification profile of *B. subtilis* rRNA was characterized, but most of the enzymes responsible for these modifications have yet to be identified (Popova et al., 2024). Prior to this, complete rRNA modification profiles had not been constructed for any Gram-positive organism, though partial profiles and individual modifications have been identified for a few Gram-positive species (Antoine et al., 2021; Emmerechts et al., 2007; Hansen et al., 2002; Jiang et al., 2018; Kirpekar et al., 2018; Maus et al., 2005).

In this work, we characterize the dual-substrate 16S and 23S 2′-O-methylcytidine (Cm) methyltransferase TlyA, previously named YqxC, in *B. subtilis*. This enzyme was renamed TlyA in *B. subtilis* (TlyA_*Bs*_) after TlyA in *Mycobacterium tuberculosis* (TlyA_*Mtb*_) (Johansen et al., 2006). TlyA_*Mtb*_ functions equivalently as a 16S/23S 2′-O-methylcytidine methyltransferase and modifies at orthologous positions on the *M. tuberculosis* rRNAs though it does not follow rRNA methyltransferase naming conventions because it was originally discovered as a hemolysin (Johansen et al., 2006; Maus et al., 2005; Wren et al., 1998). Out of the known *B. subtilis* enzymes, only RlmP and TlyA have been shown to be 2′-O-methyltransferases, and RlmP does not have any known distinct deletion phenotypes (Roovers et al., 2022). *E. coli* contains four 2′-O-methyltransferases including three that modify the 23S as Gm2251, Cm2498, and Um2552, conferred by RlmB, RlmM, and RlmE, respectively (Lövgren & Wikström, 2001; Purta et al., 2009; Widerak et al., 2005). Only RlmE mutants demonstrate a delay in 50S ribosome assembly and compromised translation initiation and elongation (Ishiguro et al., 2019; Widerak et al., 2005); RlmB and RlmM show slight reductions in fitness, but the contribution of methylation to ribosome assembly is unclear (Lövgren & Wikström, 2001; Purta et al., 2009). On the *E. coli* 16S rRNA, C1402 is dimethylated by RsmI and RsmH forming Cm and m4C modifications, respectively, to shape the P site of the ribosome and assist in mRNA decoding accuracy (Zhao et al., 2014). Therefore, Nm modifications have been shown to be important for ribosome assembly and translational fidelity in *E. coli*, but their contribution in *B. subtilis* remains unknown.

We describe the biological consequences of *tlyA*, finding that *ΔtlyA* cells exhibit a cold sensitivity phenotype and aminoglycoside antibiotic resistance. Cold sensitivity can be attributed to a ribosome assembly defect, as shown by increased levels of 50S large subunits in *ΔtlyA* cells. We also determine the identity and location of the modifications conferred by TlyA_*Bs*_ as Cm modifications at C1949 of the 23S rRNA and C1417 of the 16S rRNA which was done concurrently with the work presented in Popova et al., 2024. Additionally, we identify the catalytic site and SAM cofactor binding domain of TlyA_*Bs*_ *in vivo*, providing insight into the mechanism used to deliver methyl groups to two cytosine residues in the 16S and 23S rRNA. Importantly, we note that deletion of TlyA_*Bs*_ cannot be complemented with its ortholog from *M. tuberculosis* suggesting differing enzyme structure or substrate specificity between the two species. Taken together, this work characterizes the first and only identified rRNA methyltransferase in *B. subtilis* that modifies both the 16S and 23S rRNAs. Our work also highlights a key evolutionary difference between Gram-positive and Gram-negative bacteria, as TlyA does not have any identified homologs in any Gram-negative bacterial species.

## RESULTS

### *ΔtlyA* causes growth sensitivities and antibiotic resistance

To determine the importance of TlyA to the cell and understand its function, the *tlyA* gene was deleted in the WT (PY79) background leaving only a *loxP* scar (see MATERIALS AND METHODS). Growth of *ΔtlyA* cells was then compared to WT using spot titer assays. Ribosome assembly defects have been shown to be exacerbated by temperatures (Connolly et al., 2008). Because TlyA was hypothesized to be an rRNA methyltransferase based on sequence similarity, we first compared its growth at 37°C and 25°C (**Figure 1A**). We noticed a major growth sensitivity to cold in the *ΔtlyA* strain initially indicating its function is related to the ribosome. We normally grow *B. subtilis* at 30°C, but because of the cold sensitivity all experiments were performed at 37°C where WT and *ΔtlyA* exhibit more similar growth rates. This sensitivity was shown to be due to the loss of TlyA as full complementation was shown in *ΔtlyA* when *tlyA* was expressed from the ectopic locus *amyE* with low expression from the uninduced promoter P_hyperspank_ or overexpression with β-D-1-thiogalactopyranoside (IPTG) induction (**Figure 1A**). Overexpression of *tlyA* in a WT background did not cause a visible difference in growth rate compared to the WT control and complement in the deletion strain (**Figure 1A**).

**Figure 1.**
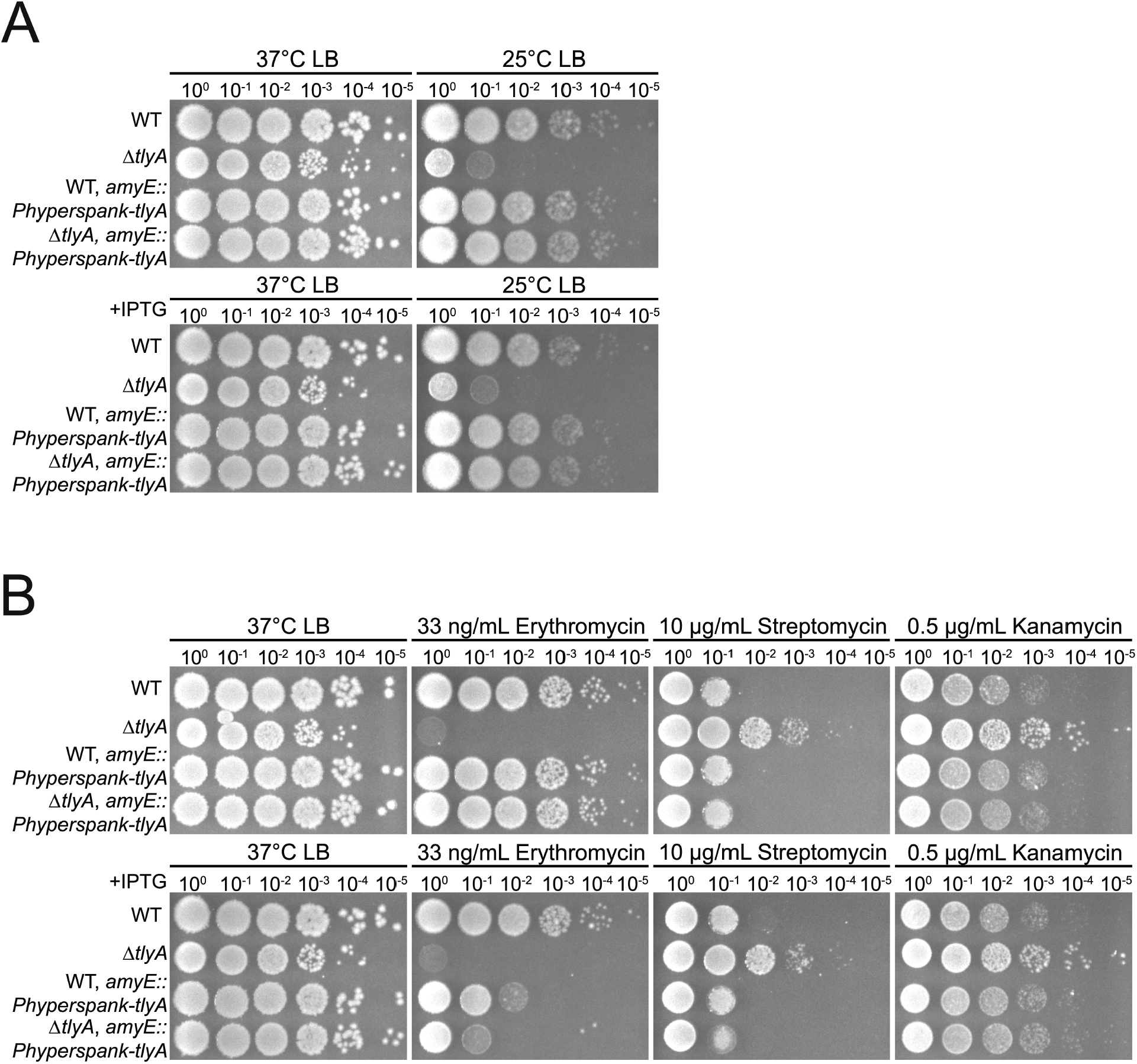
Spot-titer growth assays show cells with *ΔtlyA* are sensitive to cold and erythromycin while resistant to aminoglycosides. Shown are spot-titer dilutions of the indicated strains plated on media under the conditions listed. **(A)** Tests cold sensitivity at 25°C. **(B)** Cells were plated in the presence of the macrolide erythromycin and aminoglycosides kanamycin and streptomycin. In each case *ΔtlyA* is complemented using leak expression from the P_hyperspank_ promoter from an ectopic locus.

To determine if methylation from *tlyA* resulted in altered sensitivity to ribosome-targeting antibiotics we tested growth on plates with a variety of antibiotics including erythromycin, a macrolide targeting 23S rRNA near the peptide exit tunnel, and streptomycin and kanamycin, aminoglycosides targeting 16S rRNA near the ribosomal decoding site (Doi et al., 2016; Vázquez-Laslop & Mankin, 2018) (**Figure 1B**). Results show growth sensitivity to erythromycin and resistance to both aminoglycosides suggesting that the modification is on the 16S rRNA. These results differ from those found with the loss of TlyA in *M. tuberculosis* where *ΔtlyA*_Mtb_ shows resistance to cyclic peptide antibiotics capreomycin and viomycin but does not show cross resistance with other aminoglycosides (Johansen et al., 2006). Interestingly, the overexpression of *tlyA* with the addition of IPTG only partially rescues the growth defect when challenged with erythromycin. As shown and described in greater detail below using western blots, native expression of TlyA is similar to uninduced expression from an IPTG-regulated promoter. This result explains how uninduced expression levels, but not overexpression, of TlyA can fully complement the erm sensitivity **(Figure 5)**. Taken together, the phenotypic growth assay results demonstrate effects on antibiotic efficacy and suggest a ribosome assembly defect may occur in the absence of the Cm modifications conferred by TlyA.

### *B. subtilis tlyA* forms 2′-O-methylcytidine modifications at 23S C1949 and 16S C1417 rRNAs

Cold sensitivity and resistance to ribosome targeting antibiotics are consistent with TlyA function as an rRNA methyltransferase. Therefore, we sought to identify the modifications conferred by TlyA on the 23S or 16S rRNAs. To do this, ribosomal RNAs were isolated and analyzed by liquid chromatography coupled to tandem mass spectrometry (LC-MS/MS) (Jones et al., 2023) across three strains WT, *ΔtlyA*, and *ΔtlyA, amyE::*P_hyperspank_*-tlyA* as a complement control. The levels of 50 modifications were simultaneously quantified, including all modification types known to be conferred by *E. coli* rRNA methyltransferases (Pletnev et al., 2020). Compared to WT, *ΔtlyA* showed a significant decrease in levels of Cm modifications on both the 23S **(Figure 2A and supplemental Table S1)** and 16S **(Figure 2B)** rRNAs. Ectopic chromosomal expression of *tlyA* from the IPTG-inducible P_hyperspank_ promoter in the *ΔtlyA* background restored Cm modification of the 23S and 16S RNA. Our findings support a recent report showing that *B. subtilis* TlyA confers Cm modifications on the 23S and 16S rRNA (Popova et al., 2024). We detected very low levels of five further varieties of nucleoside methylations in the 23S and eight in the 16S **(Figure 2 and supplemental Table S1)**. Notably, these observations are unlikely to arise from tRNA contamination as none of the common tRNA specific modifications (i.e. t^6^A, i^6^A) are detected. Instead, these modifications most likely originate from trace levels of contaminating rRNAs and are detected due to the highly sensitive nature of the triple quadrupole mass spectrometry assay employed here. However, the identification of the modification by TlyA remains clear as Cm modifications are the only modification decreased in *ΔtlyA* and return to WT levels in the complementation strain.

**Figure 2.**
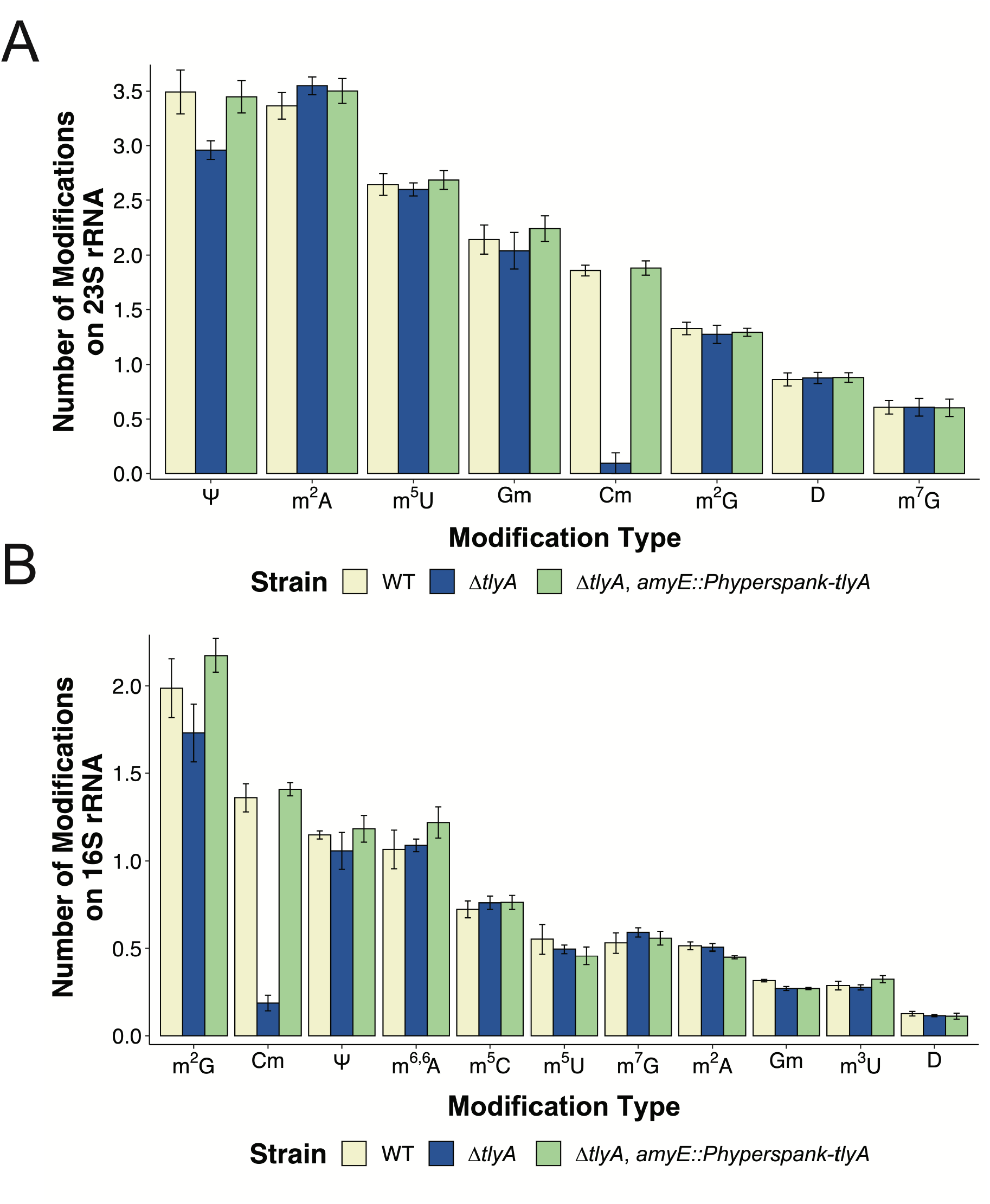
2′-O-methylcytidine modifications are absent from the 23S and 16S rRNA in *ΔtlyA*. LC-MS/MS was used to identify rRNA modifications present in WT, *ΔtlyA* and the complementing control strain. **(A)** Shows modifications detected in 23S rRNA. **(B)** Shows modifications detected in the 16S rRNA. The modifications detected are as shown N^2^-methylguanosine (m^2^G), 2′-O-methylcytidine (Cm), pseudouridine (Ψ), C2-methyladenosine (m_2_A), 5-methylcytosine (m^5^C), 5-methyluridine (m^5^U), N7-methylguanosine (m^7^G), C2-methyladenosine (m^2^A), 2′-O-methylguanosine (Gm), 3-methyluridine (m^3^U).

Given we find that *B. subtilis* TlyA is responsible for Cm modifications on the 23S and 16S rRNA, we sought to determine their location using Oxford Nanopore Technologies direct RNA sequencing. The Nanopore device pulls 5-nucleotide kmers through its pore outputting unique ionic currents and dwell times (Leger et al., 2021). Outputs differ for modified bases when compared to non-modified bases, enabling the detection of a single differentially modified base in WT and the methyltransferase deletion strain. Comparison between WT and *ΔtlyA* signal intensities and dwell times by Nanocompore analysis is represented by plotting the logistic regression log odd ratios and p-values of individual kmers to depict the likelihood of a modification between *ΔtlyA* or WT rRNAs (Johansen et al., 2006; Leger et al., 2021). Kmers in the 23S and 16S rRNAs with the lowest p-value and highest log odds ratio indicate the modified cytidine residues (Leger et al., 2021; Mulroney et al., 2023) **(Figure 3A and B)**. These results show modifications present at C1949 in the 23S rRNA and C1417 in the 16S rRNA, which are positions equivalent to those modified by *Mtb* TlyA, and support the locations recently identified for *B. subtilis* TlyA (Johansen et al., 2006; Popova et al., 2024) **(Figure 3A and B)**. Peaks generated from comparing the signal intensities and dwell times between WT and *ΔtlyA* with logistic regression log odds ratio and Kolmogorov-Smirnov tests were also generated **(Supplemental Figure 1)**. Together our LC-MS/MS data, Nanopore data, and a previous publication (Popova et al., 2024) enable us to conclude that TlyA is necessary for the formation of Cm modifications on 23S C1949 and 16S C1417 in *B. subtilis*.

**Figure 3.**
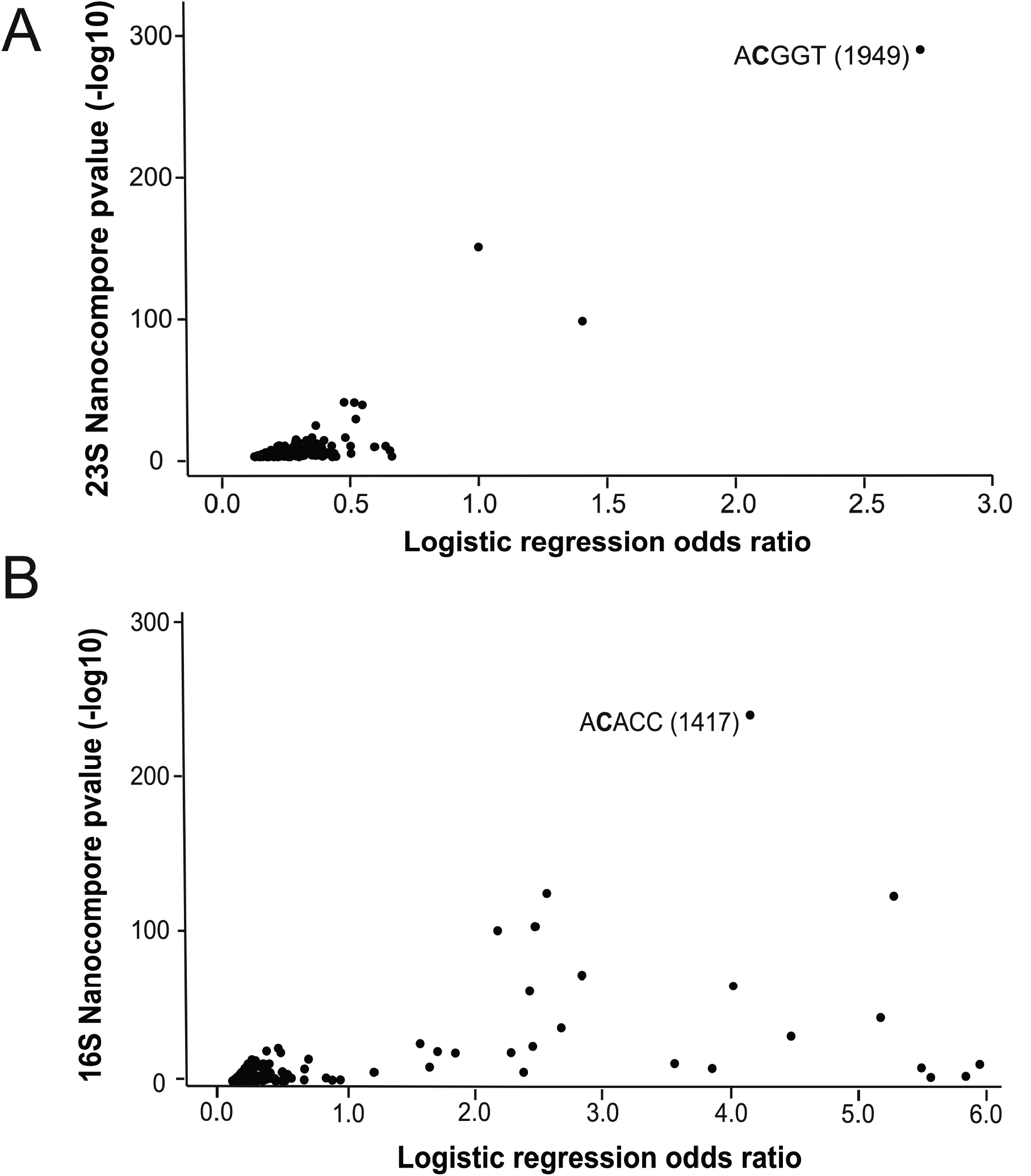
Nanopore sequencing shows modifications at C1949 in the 23S and C1417 of the 16S rRNA. Scatterplots show results of Nanocompore analyses comparing WT and *ΔtlyA* reads from **(A)** 23S and **(B)** 16S reads. The plots show the absolute value of the Nanocompore logistic regression odd ratio (GMM logit method) on the x-axis plotted against the -log10(p-value). Each point represents a p-value for a specific kmer for kmers with p-values < 0.001.

#### tlyA_Mtb_ does not complement B. subtilis ΔtlyA

Since *B. subtilis* TlyA modifies equivalent rRNA positions compared to *tlyA*_*Mtb*_ (Johansen et al., 2006), we asked if the *tlyA*_*Mtb*_ could complement *ΔtlyA* in *B. subtilis*. Though these enzymes modify equivalently in their respective species, the enzymes share only 39% sequence identity and 54% sequence similarity **(Figure 4A)**. Further, we generated an AlphaFold model of TlyA_*Bs*_ in green superimposed on the structure of TlyA_*Mtb*_ in blue, revealing many shared interior *α*– helices but visible differences in their overall structures **(Figure 4A)**. To determine if *tlyA*_*Mtb*_ can complement *B. subtilis tlyA* (*tlyA*_Bs_), we ectopically expressed *tlyA*_*Mtb*_ from *amyE* in *B. subtilis ΔtlyA* cells (see MATERIALS AND METHODS). As before, we show that *ΔtlyA*_*Bs*_ confers reduced growth at 25°C. Expression of *tlyA*_*Mtb*_ failed to complement growth of *ΔtlyA*_*Bs*_ using uninduced expression or following overexpression **(Figure 4B)**. The lack of complementation may be caused by structural differences between *B. subtilis*, and *Mtb* TlyA or differences in substrate recognition between the two enzymes. The presence of Cm modifications at homologous sites on the 23S and 16S rRNAs in *B. subtilis* and *Mtb* suggest conservation between Firmicutes, Actinomycetota and other TlyA containing bacteria. However, we find that *Mtb* TlyA is not interchangeable with *B. subtilis* TlyA and we find that *ΔtlyA*_*Bs*_ confers resistance to aminoglycoside antibiotics whereas *ΔtlyA*_*Mtb*_ does not (Johansen et al., 2006). Therefore, although *B. subtilis* and *Mtb tlyA* modify equivalent positions in the 16S and 23S, the gene deletions yield different phenotypes and *tlyA*_Mtb_ is unable to complement *ΔtlyA*_*Bs*._ These experiments suggest important differences between TlyA from *Mtb* and *B. subtilis*.

**Figure 4.**
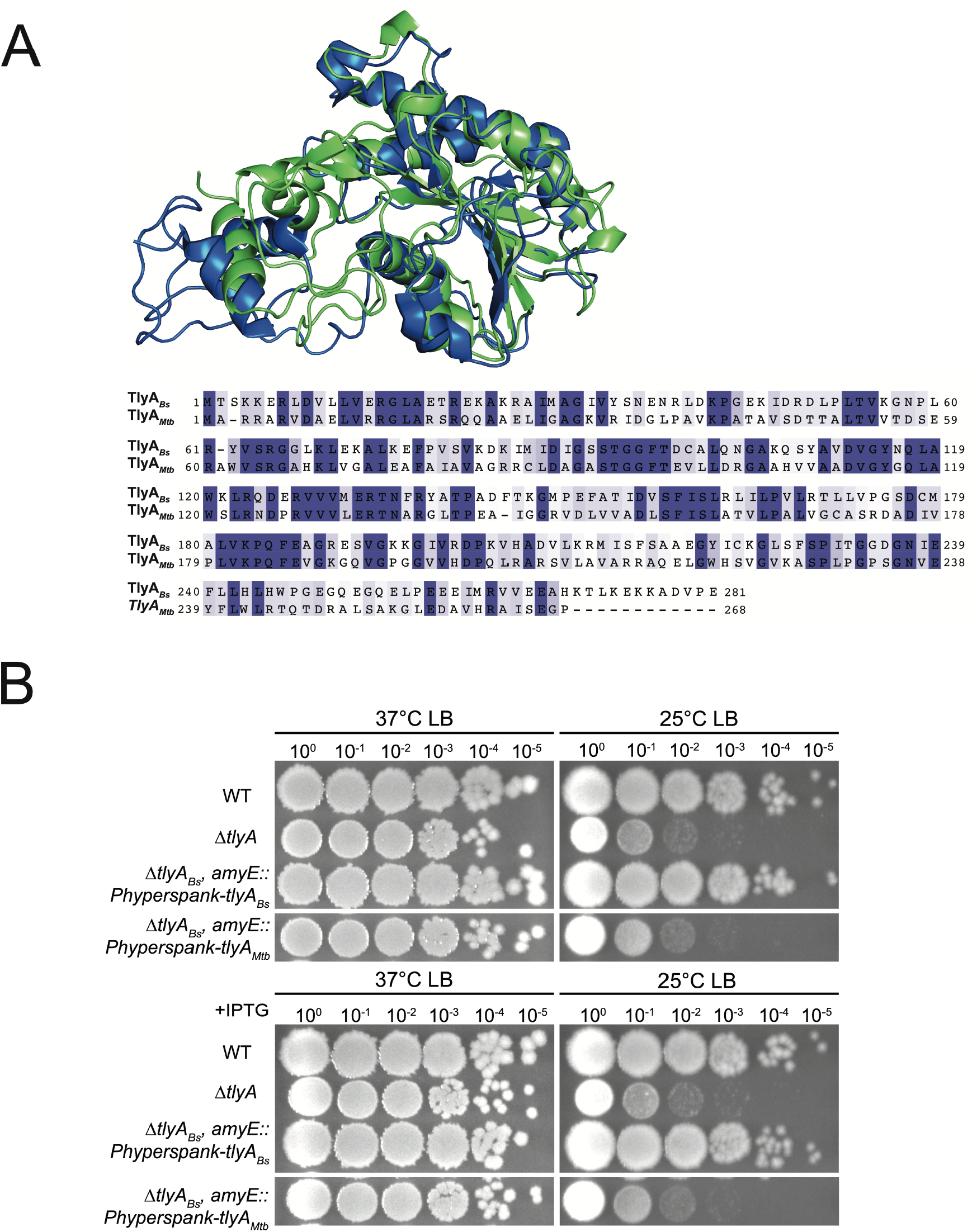
TlyA_*Mtb*_ does not complement *B. subtilis ΔtlyA*. **(A)** AlphaFold 2.0 rendition of *B. subtilis* TlyA (green) shows similarity to the structure of TlyA_*Mtb*_ (blue). Protein sequence alignment shows identical amino acids (purple) and similar amino acids (light purple) with 39% sequence identity and 54% sequence similarity shared between the two proteins. **(B)** Expression of *tlyA*_*Mtb*_ from the *amyE* ectopic locus using leak expression from an IPTG-inducible promoter fails to complement ***Δ****tlyA*.

#### Residues K69, D155, and K183 are required for catalytic activity of TlyA *in vivo*

To further understand TlyA function we investigated putative catalytic residues. The catalytic site of *B. subtilis TlyA* was proposed based on homology to the *E. coli* heat-shock methyltransferase RrmJ, as described (Hager et al., 2002). To determine residues required for *in vivo* function of *B. subtilis* TlyA, we created four mutants using site-directed mutagenesis at residues *K69, D155, K183*, and *E239* which are predicted to be important for catalysis (Hager et al., 2002). Western blotting of TlyA showed similar WT expression levels compared to TlyA variant expression **(Figure 5A, Supplementary S2)**. When expressed from an ectopic locus in a *ΔtlyA* background, the *K183A, K69A*, and *D155A* mutants were unable to rescue the phenotype of *ΔtlyA* under cold growth conditions, regardless of expression level **(Figure 5C)**. The mutant *E239A* was able to rescue the phenotype of *ΔtlyA* when induced with IPTG, indicating that overexpression of this variant was able to compensate for *ΔtlyA*. As shown in the Figure 5B model, the glutamate at position 239 is the most removed from the catalytic tetrad which could explain how only overexpression allows for complementation of the deletion. Based on the model, the K-D-K residues of the catalytic tetrad are directly stabilizing or acting on the cytosine or SAM cofactor **(Figure 5B)**. We generated a PyMOL model that depicts the SAM cofactor positioned in the active site of TlyA, where the K183 is oriented for the deprotonation of the 2′-hydroxyl and nucleophilic attack to transfer the methyl group from SAM to the cytosine. When growing our strains we noticed that the overexpression of the *K183A* mutant in *ΔtlyA* halted cell growth in LB media at 37°C. To determine if *K183A* is dominant negative to WT, we expressed *tlyA*^*K183A*^ from the P_hyperspank_ promoter in a WT background. Growth curves show that induced overexpression of *tlyA*^*K183A*^ drastically reduces cell growth during exponential phase in a WT background, suggesting this variant is indeed dominant to the WT protein **(Supplemental Figure S3)**. Given the importance of the three residues that are predicted to work directly on the substrate and cofactor, we propose that TlyA functions with the K-D-K catalytic triad, similar to RrmJ, with the glutamate supporting catalytic activity (Hager et al., 2002).

**Figure 5.**
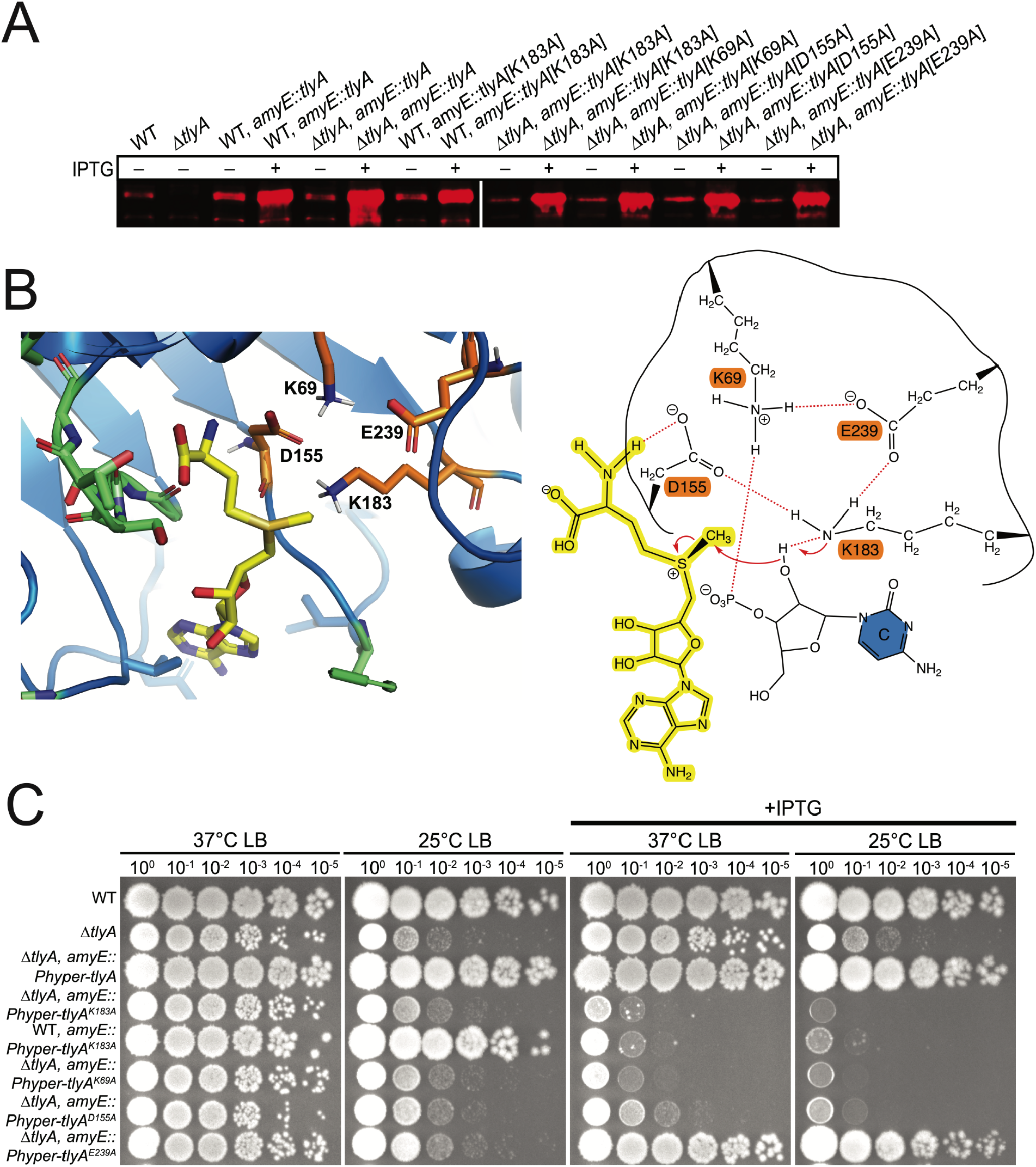
Residues K99, D155, K183 and E239 are important for function of TlyA in vivo. **(A)** Western blot of TlyA shows mutants are expressed and accumulate to WT levels and are expressed with and without addition of IPTG. **(B)** PyMOL model showing SAM cofactor positioning in the catalytic tetrad of TlyA. Catalytic residues are colored in orange with the supporting SAM-binding residues shown in green. ChemDraw illustration of the proposed methyl transfer reaction of TlyA with the SAM cofactor shown in yellow, catalytic residues in orange and the cytidine bearing the 2′-hydroxyl substrate in blue. **(C)** Spot titer assays showing failure of TlyA variants to complement the growth defect of ***Δ****tlyA* expressed from the ectopic locus *amyE* from an IPTG inducible promoter.

### B. subtilis TlyA uses the GxSxG motif to bind SAM in vivo

After determining residues important for *in vivo* catalytic function, we asked which residues could be important for TlyA to bind the S-adenosyl-L-methionine (SAM) cofactor. Many rRNA methyltransferases contain Class I MTase structure with the GxGxG motif, widely recognized to bind SAM (Kagan & Clarke, 1994; Schubert et al., 2003). *Mtb* and *B. subtilis* TlyA both contain ^90^GxSxG^94^, a variant of the GxGxG motif (Witek et al., 2017) **(Figure 6B)**. In addition to the GxSxG motif, TlyA_*Mtb*_ also requires the novel motif ^60^RXWV^63^ in the NTD to CTD linker (Witek et al., 2017). The RXWV motif organizes the GxSxG motif with the W62 and V63 being the most important for SAM affinity (Witek et al., 2017). The corresponding residues in *B. subtilis* TlyA, ^60^LRYV^63^, show the conserved valine in the linker region. To determine if these residues are important for TlyA function *in vivo* we again utilized *tlyA* mutant complementation growth assays. Western blotting shows that all TlyA variants are expressed at similar levels to WT **(Figure 6A, Supplementary S2)**. Spot titer assays demonstrate that substitution of one glycine can be tolerated, but the substitution of both glycine residues results in the inability of ectopically expressed TlyA variants to rescue the *ΔtlyA* phenotype at 25°C **(Figure 6C)**. When just a single glycine was substituted for a negatively charged residue (Glu), TlyA was nonfunctional *in vivo*. Upon mutating V63 to alanine, no growth defects were observed, demonstrating that TlyA_*Bs*_ does not require the same linker region as TlyA_*Mtb*_ does during SAM binding **(Figure 6C)**. Together, our results support the conclusion that TlyA_*Bs*_ binds SAM via its ^90^GxSxG^94^ motif *in vivo*. The linker region, with its conserved valine residue, is not required for SAM binding by *B. subtilis* TlyA *in vivo*, demonstrating yet another functional difference between *B. subtilis* and the *Mtb* TlyA enzymes.

**Figure 6.**
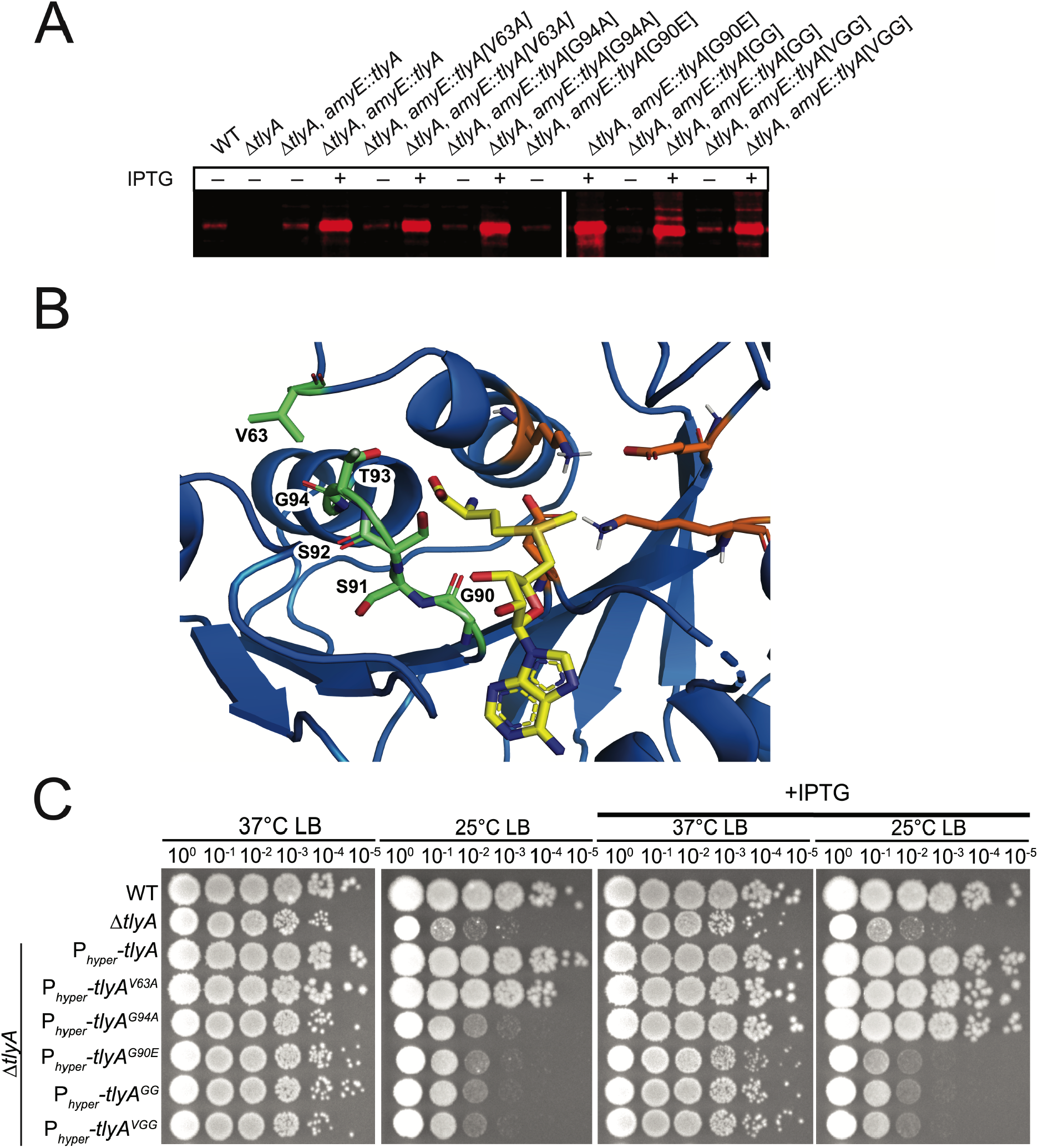
The GxSxG motif helps stabilize the SAM cofactor in the active site of TlyA *in vivo*. **(A)** Western blot shows SAM-binding mutants are expressed and accumulate to native TlyA levels *in vivo*. **(B)** AlphaFold 2.0 depiction of TlyA_*Bs*_ (blue) with residues in green (G90-G94 and V63) used to help stabilize the binding of SAM (yellow). **(C)** Spot-titer assay shows *tlyA* mutant complementation of ***Δ****tlyA* growth to at 25°C to identify mutants that are important of SAM binding *in vivo* as judged by failed complementation. GG corresponds to *G90A, G94A* and *VGG* corresponds to *V63A, G90A, G94A* all of which were expressed from *amyE* using the P_hyperspank_ promoter in a ***Δ****tlyA* genetic background where indicated.

#### Loss of *tlyA* causes an accumulation of the large subunit in associative conditions

Since TlyA from *Mtb* and *B. subtilis* methylate the same sites yet appear to have important differences and because *ΔtlyA* cells showed a growth defect at 25°C, we investigated the possible contribution of *tlyA* to ribosome assembly. To test ribosome assembly, we used sucrose gradient ultracentrifugation to quantify the formation of ribosomal subunits to determine if ribosome assembly defects resulted from the absence of TlyA. We grew WT and *ΔtlyA* cultures at both 37°C and 25°C in associating conditions where the ribosomal subunits come together to form the 70S mature ribosome and dissociating conditions where they remain as individual 50S and 30S subunits. Results from cultures grown at 37°C show no major differences in dissociating conditions but do show a decrease in mature 70S formation in associating conditions **(Figure 7A)**. This difference is further exacerbated by cold growth conditions at 25°C, where the mature 70S peak is reduced in *ΔtlyA* compared to WT and there is a significantly higher level of 50S in *ΔtlyA* compared to WT **(Figure 7B,C)**. No major differences were observed between WT and *ΔtlyA* under dissociating conditions when comparing the 50S and 30S peaks at either temperature. With these results, we show that one or both Cm modifications are necessary for the proper maturation of the 70S ribosome, and that ribosome maturation is adversely affected in the absence of TlyA especially at 25°C, causing an accumulation of the large subunit.

**Figure 7.**
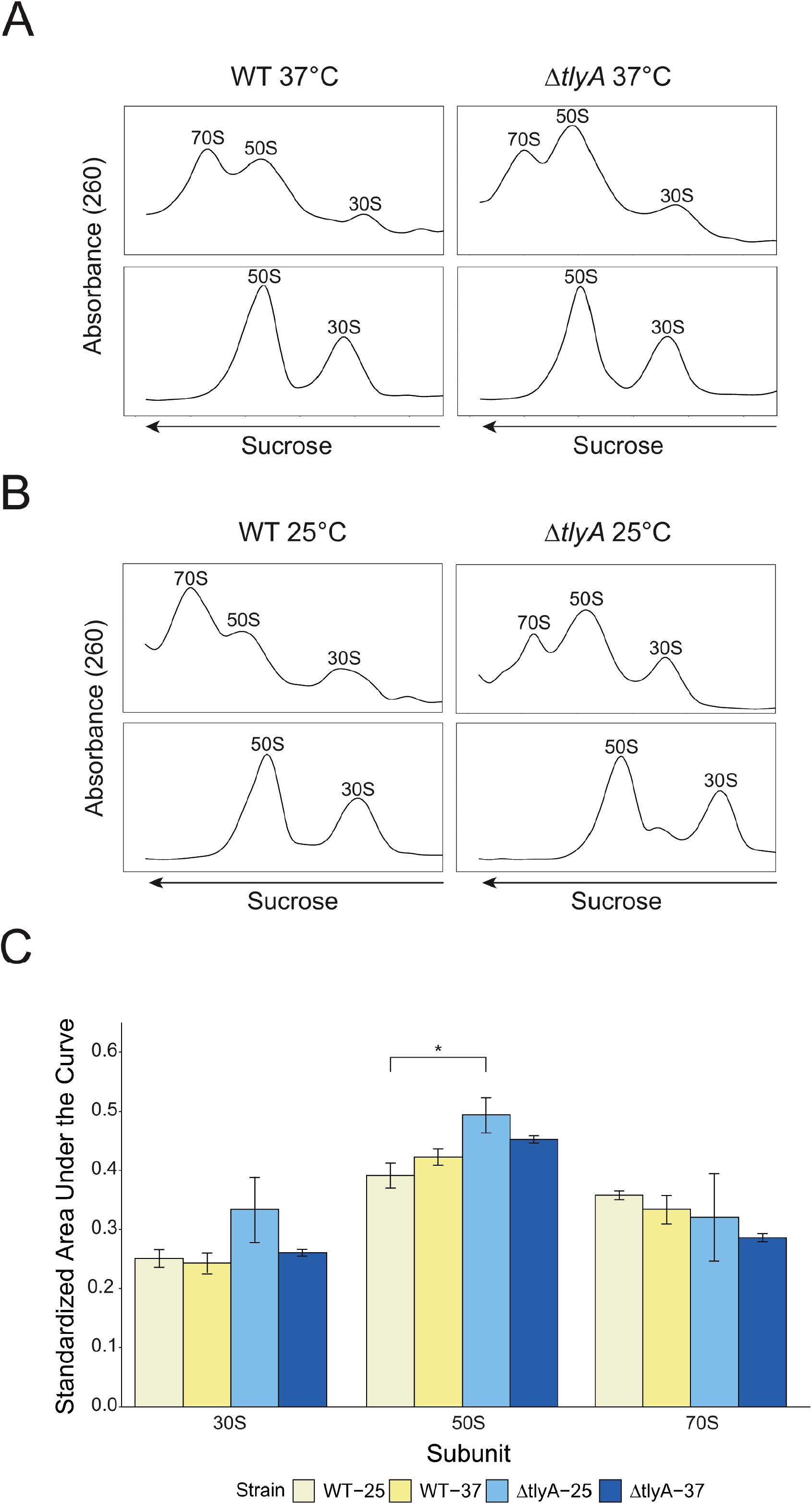
70S ribosome assembly is reduced in *ΔtlyA* cells. Fractions from sucrose gradient ultracentrifugation using associative (top panels) or dissociative bottom panels for WT and *ΔtlyA* cells grown at **(A)** 37°C or **(B)** 25°C. **(C)** Associating sucrose gradient quantification of mean area under the curve for each subunit. Black bars represent standard error, and asterisk denotes significance (* p < 0.05).

## DISCUSSION

In this study we characterize the *tlyA* gene (previously *yqxC*), as a dual-substrate 23S and 16S 2′-O-methylcytidine (Cm) rRNA methyltransferase important for ribosome assembly in *B. subtilis*. Upon deletion of *tlyA*, cell growth is perturbed at cold temperatures and when challenged with the macrolide erythromycin, an antibiotic targeting the large ribosomal subunit. Deletion of *tlyA* also confers resistance to aminoglycosides streptomycin and kanamycin, which target the small ribosomal subunit a result different from loss of *tlyA* in *Mtb* which only shows resistance to capreomycin and viomycin (Johansen et al., 2006). We also show that *tlyA* is important for assembly of mature 70S ribosomes. Prior work has identified some of the genes encoding rRNA methyltransferases in *B. subtilis*; however, none have been shown to have distinct phenotypes that effect growth or ribosome assembly (Desmolaize et al., 2011; Nishimura et al., 2007; Roovers et al., 2022; Wolff et al., 2024). Hence, our study of TlyA is uniquely important for understanding ribosome biogenesis and function in Gram-positive and other related bacteria.

We found the phenotypes of *ΔtlyA* are due to loss of 2′-O-methylcytidine modifications on the 23S and 16S rRNA and maintenance of all other methylation modifications present on rRNA as shown by our LC-MS/MS results. Our results generally match recent LC-MS/MS data for *B. subtilis* 23S and 16S rRNAs which was published as our work was ongoing (Popova et al., 2024). Utilizing Oxford Nanopore direct RNA sequencing, we determined the location of these modifications to be 23S C1949 and 16S C1417. In the structure of the ribosome these modifications exist on helix 69 of the 23S rRNA and helix 44 of the 16S rRNA. Helix 44 is part of the aminoacyl (A) site that accepts incoming tRNAs, and helix 69 is located between the A and P sites and has been shown to be essential for decoding mRNA and bridging to helix 44 of the 16S (Kulik et al., 2017; Tsai et al., 2013). Biochemically, the formation of Cm modifications by TlyA would be expected to reduce the reactivity of two different 2′-hydroxyl groups, which increases RNA stability and promotes correct conformation of the rRNA suggesting these modifications occur in functionally important areas of the ribosome needed for ribosome formation and function.

Homology search of *B. subtilis* TlyA identifies the dual substrate rRNA methyltransferase TlyA from *Mtb*. TlyA_*Mtb*_ confers Cm modifications at homologous rRNA sites that we report for *B. subtilis* (Johansen et al., 2006). We were interested in the possibility that TlyA function is conserved across species, since we did not detect similar TlyA enzymes in *E. coli* or other more divergent Gram-negative species. We found that *ΔtlyA*_*Bs*_ could not be complemented by *tlyA*_*Mtb*_, despite their similarity, suggesting that structural differences may exist between these methyltransferases. Further, *ΔtlyA*_*Bs*_ confers resistance to aminoglycosides whereas *ΔtlyA*_*Mtb*_ does not (Johansen et al., 2006). Therefore, differences between these enzymes could be important for their ability to recognize and modify native rRNA sequences prohibiting cross-species complementation and differences in native deletion phenotypes. We wish to note that there are no known rRNA methyltransferases in Gram-negative bacteria that modify both the large and small ribosomal subunits. Therefore, the presence of the dual substrate rRNA methyltransferase TlyA highlights a distinction in ribosome maturation between Gram-positive (Bacillus), diderm (Mycobacterium) and Gram-negative bacteria.

To identify residues important for catalytic function, we identified residues in the catalytic domain of TlyA_*Bs*_ conserved with the Gram-negative rRNA methyltransferase RrmJ/RlmE. The active site housed in the CTD of TlyA_*Bs*_ contains a RrmJ/FtsJ Rossmann-like methyltransferase fold. As previously mentioned, RrmJ is a heat shock-induced 23S rRNA methyltransferase in *E. coli* and uses the catalytic tetrad of residues K38, D124, K164, and E199 in the methyl transfer reaction (Hager et al., 2002). When we mapped residues K69, D155, K183, and E239 to TlyA_*Bs*_, we hypothesized that these residues are positioned to catalyze the methyl transfer in the active site. Residues K69, D155, and K183 are indispensable for the function of TlyA in *B. subtilis*; however, TlyA^E239^ can only complement *ΔtlyA* when overexpressed from an ectopic locus. Defining the residues necessary for the *in vivo* function of TlyA_*Bs*_ demonstrates that active site residues are similar between rRNA methyltransferases from Gram-positive and Gram-negative species.

TlyA_*Mtb*_ was used to determine the conserved S-adenosyl-L-methionine (SAM) binding residues used in TlyA_*Bs*_. *In vitro* assays demonstrate that TlyA_*Mtb*_ utilizes the SAM binding motif I GxSxG and a novel RXWV tetrapeptide motif linking the N-terminal domain (NTD) and C-terminal domain (CTD) to hold SAM in the active site (Laughlin & Conn, 2019; Witek et al., 2017). TlyA_*Bs*_ has residues G90, S91, S92, T93, and G94, comprising the GxSxG motif, and V63 in the linker region. We tested residues that are required for SAM binding in TlyA_*Mtb*_ and are present in TlyA_*Bs*_ to elucidate the degree of enzymatic conservation between orthologs. In TlyA_*Bs*_, we found the glycine residues in the SAM motif I but not the valine linker is necessary for function *in vivo*. Our results illustrate that for *B. subtilis*, TlyA only requires the GxSxG motif *in vivo*, indicating that there may be functional differences in TlyA rRNA methyltransferases across bacteria. However, our results are limited in their *in vivo* study because cell growth was used as an indirect measure of TlyA variant function. The residues we find to be required for *in vivo* SAM binding in *B. subtilis* highlight an enzymatic difference between TlyA_*Bs*_ and TlyA_*Mtb*_ that may prevent the enzymes from functioning interchangeably.

Our work identifies the first rRNA methyltransferase in *B. subtilis* that shows a distinct phenotype including a ribosome assembly defect. An rRNA methyltransferase modifying both the 23S and 16S is not present in *E. coli* and other Gram-negative bacteria demonstrating an evolutionary difference between Gram-positive, diderm and Gram-negative bacteria. It will be important to learn if other rRNA methyltransferases in *B. subtilis* have similar phenotypes and effects on ribosome assembly, or if TlyA is the most crucial rRNA methyltransferase for ribosome biogenesis.

## MATERIALS AND METHODS

### Strains and media

Bacterial strains were all derived from *Bacillus subtilis* lab strain PY79. Strains used, primers, and plasmids are listed in supplement tables S2-S4. *B. subtilis* strains were regularly grown in LB broth (10 g/L tryptone, 10 g/L NaCl, 5 g/L yeast extract). LB agar used to plate strains used the same ratio but with 15 g of agar supplemented. Plates and cultures were grown regularly at 37°C instead of 30°C to mitigate the cold defect seen in *ΔtlyA*. When necessary, broth or plates were supplemented for 100 µg/mL spectinomycin or 0.5 µg/mL erythromycin for selection. Freezer stocks of bacterial strains were composed of 900 µL of culture grown to OD600 > 1.0 and 100 µL dimethyl sulfoxide (DMSO) and stored at -80°C.

### ΔtlyA strain construction and transformation

*B. subtilis yqxC::loxP-erm-loxP trpC2* strain was obtained from the Bacillus Genetic Stock Center (BGSC) (Koo et al., 2017). Genomic DNA was isolated from this strain and transformed into competent WT cells. To make competent, WT bacteria was inoculated into LB supplemented with 3 mM MgSO_4_ and grown on a rolling rack at 37°C for ∼3 hours or until reaching an OD600 of >1.0. Cultures were back diluted 1:25 into 500 µL MD media (1x PC buffer [10x PC buffer: 107 g/L K_2_HPO_4_, 60 g/L KH_2_PO_4_, 10 g/L trisodium citrate, (H_2_O)5], 2% glucose, 50 µg/mL tryptophan, 50 µg/mL phenylalanine, 11 µg/mL ferric ammonium citrate, 2.5 µg/mL potassium aspartate, 3 mM MgSO_4_) and grown for 4 hours on a rolling rack at 37°C. No more than 500 ng DNA was added to the competent culture and transformed for 1-1.5 hours on a rolling rack at 37°C. Transformation was confirmed using antibiotic selection and PCR. Subsequent clean deletion made us of the Cre recombinase expressing plasmid pDR244 as described previously (Wozniak & Simmons, 2022).

### ΔtlyA complementation

For genetic complementation, we cloned *tlyA* or mutant *tlyA* into the modified version of plasmid pDR110 (pPB194) containing the P_hyperspank_ promoter and spectinomycin selection cassette flanked by *amyE* regions. To do this, our gene of interest was amplified with primers containing overhanging complementary regions to our plasmid. These were combined with amplified plasmid backbone and Gibson assembled according to manufacturer’s recommendations (NEB E2611). The Gibson reaction was then transformed into MC1061 *E. coli*, successful transformants selected by ampicillin, and mini-prepped according to manufacturer’s instructions (Qiagen 27104). Miniprepped plasmid sequences were confirmed with Sanger sequencing. Following, *ΔtlyA B. subtilis* was made competent and transformed as described above resulting in homologous recombination at the *amyE* locus. This disruption of the *amyE* gene was confirmed by the strain’s inability to metabolize starch by growth on a LB agar plate containing 10 g/L starch and then stained with solid iodine. Strains with successful recombination as indicated by failure to metabolize starch underwent PCR of the *amyE* locus to ensure correct variant length of the disrupted locus.

### Genomic DNA (gDNA) purification

Desired strains were struck out onto the appropriate LB agar plates and incubated at 37°C overnight. A single colony was used to inoculate 5 mL LB media, shaking at 200 rpm at 37°C for 3 hours, or until OD600 reaches 2.0. Cells were pelleted at 8,000 x g for 5 minutes at room temperature and the pellet was resuspended in 200 µL lysis buffer (50 mM Tris pH 8.0, 10 mM EDTA, 1% Triton X-100, 0.5 mg/mL RNase A, 20 mg/mL lysozyme) and incubated at 37°C for 30 minutes. 20 µL of 10% sodium dodecyl sulfate (SDS) and 20 µL protease K (10 mg/mL in TE buffer with 10% glycerol) were added and incubated at 55°C for 30 minutes. 500 µL of PB buffer (5 M GuHCl, 30% isopropanol) was added and the entire sample volume was loaded onto a Qiagen QIAprep Spin Miniprep column and centrifuged at 13,000 x g for 1 minute, followed by another 500 µL buffer wash. 750 µL of PE buffer (10 mM Tris pH 7.5, 80% ethanol) was added and centrifuged at 13,000 x g for 30 seconds, followed by one dry spin for 1 minutes. Columns were transferred to a new microcentrifuge tube and gDNA was eluted with 100 µL ddH_2_O at 13,000 x g for 1 minute, reloaded and spun again. The gDNA concentration and purity was measured using a Nanodrop and samples were stored at -20°C.

### Spot titer sensitivity assays

*B. subtilis* strains were struck out onto the appropriate LB agar plates and incubated at 37°C overnight. A single colony per strain was inoculated in 2 mL LB media in 14 mL round bottom culture tubes, supplemented with spectinomycin when necessary. Cultures were incubated on a rolling rack at 37°C until the OD600 was 0.5-0.8. Cultures were normalized in 200 µL to an OD600 of 0.5 using 0.85% w/v sterile saline and were serially diluted in saline to 10^−5^. Plates were prepared the day of the assay with appropriate antibiotic and/or 100 µM IPTG as indicated in figures. Five µL of the serial dilutions were spotted onto prepared LB agar plates. Plates were grown at 37°C for 11 hours or 25°C for 36 hours. All spot titers were performed in biological triplicate and were imaged using the white light source in the Alpha Innotech MultiImage Light Cabinet, with exposure, brightness, and contrast edited in Adobe Photoshop.

### Total RNA isolation

Total RNA isolation was done for LC-MS/MS and Nanopore procedures. For LC-MS/MS, 50 mL cultures were inoculated with a single colony of appropriate strain. Antibiotics and IPTG were added to the *tlyA* complement cultures. Strains were grown until they reached an OD600 of 0.6-0.8. Forty mL were then centrifuged at max speed of tabletop centrifuge for 15 min. Pellets were then resuspended in 5.6 mL NE (0.1 M NaCl, 0.05 M EDTA) and incubated for 5 min at 37°C. Following 300 µL of freshly made lysozyme (40 mg/mL in water) was added to each tube and incubated at 37°C for 15 min. Following incubation, 700 µL 10% sarkosyl was added and incubated on ice for 5 min. Lysates then underwent phenol:chloroform extraction by adding one volume of phenol:chloroform:iso-amyl alcohol (125:24:1) (Sigma P1944), shaking and vortexing, and spinning down table top centrifuge max speed for 30 min at 4°C. The aqueous phase was pipetted off the top and underwent an additional phenol extraction, followed by a final chloroform extraction. To precipitate the nucleic acids, 3X volumes 100% ethanol was added with 0.1X volume of 3M NaOAc to the extraction, inverted to mix, and placed at -20°C for at least 2 hours up to overnight. Following precipitation, nucleic acids were pelleted at max speed for 10 min at 4°C in tabletop centrifuge, washed twice with 70% ethanol, and dried at RT for ∼10 min. The pellet was then resuspended in 5 mL water and precipitated again by mixing with 3X volumes ethanol and ½ volume 7.5M ammonium acetate and placing at -20°C for at least 2 hours up to overnight. Following second precipitation, nucleic acids were pelleted and washed again as previously described. The pellet was then resuspended in 400 µL water, transferred to a microcentrifuge tube and incubated with DNase I (Roche 04716728001) according to manufacturer’s instructions at 37°C. The reaction was quenched after 30 min with one phenol:chloroform:iso-amyl alcohol extraction and one chloroform extraction. RNA was precipitated at -20°C overnight with the addition of 3X ethanol and 0.1X NaOAc. Following, RNA was pelleted, washed with 70% ethanol, and dried as described above and resuspended in 80 µL water. Concentrations of 1:20 diluted product was determined by nanodrop for approximate or qubit (Broad Range RNA kit) for precise measurement. RNA quality was checked using bleach gel agarose electrophoresis (Aranda et al., 2012). Total RNA was then aliquoted and stored at -80°C for long-term storage. Total RNA extraction for Nanopore uses was done in a the same but was scaled down in volume starting with an initial 5 mL culture. Additionally for these isolations there was no additional phenol extraction upon initial clean up and the second pelleting, washing, and precipitation was abandoned as higher purity was not as necessary for these applications. For LC-MS/MS applications 23S and 16S rRNAs were further isolated using 1% agarose 0.5X TBE gel electrophoresis and the Zymoclean Gel RNA Recovery kit (Zymo R1011) according to manufacturer’s instructions.

### Quantitative ribonucleoside LC-MS/MS

The ribonucleoside modification landscape was quantitatively assessed using a highly multiplexed targeted ribonucleoside LC-MS/MS assay as previously described (Jones et al., 2023). Briefly, purified rRNA (150 ng) was enzymatically degraded to monoribonucleosides using a two-step digestion. The RNA was first hydrolyzed to ribonucleotide monophosphates using 300 U nuclease P1 (NEB, 100,000 U/mL) per μg of RNA overnight at 37°C in 100 mM ammonium acetate (pH 5.5) and 100 μM ZnSO_4_. The resulting ribonucleotides were dephosphorylated using 50 U bacterial alkaline phosphatase (Invitrogen, 150 U/μL) per μg of RNA for 4 hr at 37°C in 100 mM ammonium bicarbonate (pH 8.1) and 100 μM ZnSO_4_. Prior to the digestion, the enzymes were buffer exchanged into the respective digestion buffers using a BioRad Micro Bio-Spin 6 size exclusion column to remove glycerol and other ion suppressing contaminates. The resulting monoribonucleoside mixture was lyophilized and resuspended in 10 μL 40 nM ^15^N_4_-inosine as an internal standard. Following, the ribonucleosides were separated by reversed phase chromatography and quantified using multiple reaction monitoring on a triple quadrupole mass spectrometer as previously described (Jones et al., 2023).

### Nanopore

Ribosomal 23S or 16S RNAs were pulled down from total RNA of either WT or *ΔtlyA* strains using custom-ordered reverse transcription adapters as described in the Oxford Nanopore Technologies Direct RNA Sequencing Kit manual (SQK-RNA002). Samples were then generated according to kit instructions and ran with a R9.41 flow cell (FLO-MIN106) using ONT’s MinION Mk1B technology. Two biological replicates were used for WT and *ΔtlyA* for each of the 23S and 16S rRNAs. Reads from WT and *ΔtlyA* were then compared using the Nanocompore pipeline according to its online manual to indicate modification location by analyzing differences in electrical current peaks and dwell time (Leger et al., 2021).

### Site-directed Mutagenesis

Site-directed mutagenesis was carried out using the efficient one-step single-site plasmid mutagenesis protocol, as previously described (Liu & Naismith, 2008). The DNA template used for amplification was pPB41, containing a wild-type copy of *tlyA* under the IPTG-inducible P_hyperspank_ promoter. Four catalytic mutants were constructed at residues K69, D155, K183, and E239 using the primers described in supplemental table. Six SAM-binding mutants were created at residues V63, G90, G94 using primers described in supplemental table. Plasmids were transformed into competent MC1061 and mini-prepped according to manufacturer’s instructions (Qiagen 27104). All constructs were Sanger sequenced to ensure *tlyA* was correctly mutated. Plasmids were transformed in *ΔtlyA*.

### Western blot

To ensure that the functionally mutated copies of TlyA were properly expressed, we conducted Western blots using a custom polyclonal antiserum against TlyA. One colony was inoculated per strain in 25 mL LB media and was grown at 37°C for approximately two hours until reaching an OD_600_ of ∼0.5-0.8. Strains containing *tlyA* under the P_hyperspank_ promoter were either left uninduced or induced with 100 µM IPTG once at an OD_600_ between ∼0.15-0.25 as noted in the figure. Cells were spun down and resuspended in 1 mL of lysis buffer (20 mM Tris pH 8.0, 4M NaCl, 1M DTT, 0.5M EDTA, 10% SDS, urea, and water) and further lysed by sonication 60 Hz for 1 min total (10 sec on/10 sec off). Crude cell extracts were boiled for 5 min, mixed with 2X SDS loading dye, and run on SDS-PAGE. Following, proteins were transferred to Cytiva Amersham nitrocellulose membranes using the Bio-Rad Trans-blot Turbo System set for using the system’s setting mixed weight protein transfer. Following transfer, membranes were incubated in 5% milk for blocking, followed by incubation with anti-TlyA (1/500 dilution) antibody for 1 hour at room temperature. The membrane was then washed four time with TBS-T and incubated with fluorescent tagged LICOR 800 anti-Rabbit secondary antibody (Fisher 92632211) for 1 hour at room temperature. After, the membrane was rinsed four times with TBS-T and imaged with a Licor Odessey Fc Imager.

### Growth curve of catalytic mutant

*B. subtilis* strains were struck out onto the individual LB agar plates supplemented with spectinomycin agar when needed and incubated at 37°C overnight. Strains were plate washed and standardized to a starting OD_600_ of 0.02 and inoculated in 12.5 mL of LB media in 125 mL Erlenmeyer flasks. Cultures were grown shaking at 200 rpm and 37°C for a total of 7 hours, where the OD_600_ measurement was taken every 20 minutes. Since the overexpression of TlyA-K183A appeared to be catalytically lethal, IPTG was added at the beginning of the exponential phase (OD_600_ = 0.2). Growth measurements were taken in biological triplicate and were analyzed and graphically displayed using R.

### Sucrose gradient ultracentrifugation

Ribosomal subunit analysis was done by sucrose gradient ultracentrifugation as previously described (Roovers et al., 2022). *B. subtilis* cultures of 150 mL were grown up at 37°C or 25°C to an OD600 of 0.5-0.6. Cultures were pelleted via centrifugation at 4,500 x g for 10 min, and each pellet was resuspended in 150 µL buffer G (20 mM Tri-HCl pH 7.5, variable Mg(OAc)_2_, 200 mM NH_4_Cl, 6 mM β-mercaptoethanol) with either associating (10 mM) or dissociating (1 mM) Mg(OAc)_2_, 1 µL 50 mg/mL lysozyme, and 1X protease inhibitors (Pierce, EDTA-free). Resuspended cells were then lysed by sonication on ice. Following, lysates were DNase I (Roche 04716728001) treated for 10 min on ice following manufacturer’s instructions. Lysate was cleared by centrifugation in a tabletop microcentrifuge for 30 min at 16000 rpm and 4°C. Approximately 15 A_260_ units were added to the top of each sucrose gradient [10-40% (w/v) in association or dissociating buffer conditions] and were ultracentrifuged in a Thermo TH-641 rotor at 25,000 rpm for 15 hr at 4°C. Gradients were fractionated by hand into UV translucent 96 well plates. Each fraction was diluted 1:10 and the dilutions were measured for A_260_ by Tecan Infinite M100 or Tecan Spark spectrophotometers. Three biological replicates were used to quantify mean area under the curve for every strain and subunit. Subunit areas were separated based on the visualization of their peaks. Area under the curve was quantified using the Simpson’s Rule method and normalized by dividing each subunit’s area by the total area of all three. Significance was determined using an ANOVA with post hoc analysis using the Bonferroni method.

## Data availability

All Nanopore sequencing data used to identify modification location sites is available at Sequence Read Archive (SRA) by NCBI through accession number <u>PRJNA1237866</u>. All other resources are described.

## ACKNOWLEGEMENTS

We thank Dr. Miten Jain for his help in resolving issues with analyzing our Nanopore data. We also thank Keerthikka Ravi for her insights and answers to our many coding questions. Further, we thank Roberto Bloom for his initial help in the characterization of the *tlyA* deletion. JLH was supported in part by the Genetics Training Program Grant T32–GM007544. JDJ was supported in part by an NSF GRFP. This work was supported by funding from the National Institutes of Health grant R35GM131772 to LAS and R01HG013876 to KSK.

