## Supporting tables and figures for "TlyA is a 23S and 16S 2′-O-methylcytidine methyltransferase important for ribosome assembly in *Bacillus subtilis*"

**Table S2: Strains used in this work**

| Strains | Genotype | Citation |
|---|---|---|
| JLH005 | WT PY79 | |
| JLH010 | Δ*tlyA* | This work |
| JLH018 | Δ*tlyA, amyE::P*$_{hyperspank}$-*tlyA* | This work |
| JLH020 | MC1061 | |
| JLH022 | WT, *amyE::P*$_{hyperspank}$-*tlyA* | This work |
| JLH155 | Δ*tlyA, amyE::P*$_{hyperspank}$-*tlyA*[*K183A*] | This work |
| JLH056 | Δ*tlyA, amyE::P*$_{hyperspank}$-*tlyA*[*V63A*] | This work |
| JLH162 | Δ*tlyA*$_{Bs}$, *amyE::P*$_{hyperspank}$-*tlyA*$_{Mtb}$ | This work |
| JLH164 | Δ*tlyA, amyE::P*$_{hyperspank}$-*tlyA*[*K69A*] | This work |
| JLH165 | Δ*tlyA, amyE::P*$_{hyperspank}$-*tlyA*[*D155A*] | This work |
| JLH166 | Δ*tlyA, amyE::P*$_{hyperspank}$-*tlyA*[*E239A*] | This work |
| JLH167 | WT, *amyE::P*$_{hyperspank}$-*tlyA*[*K183A*] | This work |
| JLH168 | Δ*tlyA, amyE::P*$_{hyperspank}$-*tlyA*[*G94A*] | This work |
| JLH171 | Δ*tlyA, amyE::P*$_{hyperspank}$-*tlyA*[*G90E*] | This work |
| JLH172 | Δ*tlyA, amyE::P*$_{hyperspank}$-*tlyA*[*G90A+G94A*] | This work |
| JLH173 | Δ*tlyA, amyE::P*$_{hyperspank}$-*tlyA*[*V63A+G90A+G94A*] | This work |

**Table S3: Plasmids used in this work**

| Plasmid | Vector | Insert | Reference/Source |
|---|---|---|---|
| pDR244 | pDR243 | Cre-recombinase | Bacillus Genetic Stock Center (ECE274) |
| pJLH004 | pPB194[1] | *tlyA* | This study |
| pLMM005 | pPB194 | *tlyA*[*K183A*] | This study |
| pLMM006 | pPB194 | t*lyA*[*V63A*] | This study |
| pLMM014 | pPB194 | t*lyA*[*K69A*] | This study |
| pLMM015 | pPB194 | t*lyA*[*D155A*] | This study |
| pLMM016 | pPB194 | t*lyA*[*E239A*] | This study |
| pLMM017 | pPB194 | t*lyA*[*G94A*] | This study |
| pLMM018 | pPB194 | t*lyA*[*G90E*] | This study |
| pLMM019 | pPB194 | t*lyA*[*G90E+G94A*] | This study |
| pLMM020 | pPB194 | t*lyA*[*G90E+G94A+V63*] | This study |

**Table S4: Primers used in this work:** Overhangs are underlined. Mutated nucleotides are bolded for mutagenesis primers.

| Primer | Sequence | Purpose |
|---|---|---|
| PEB3F | GCTAGCC<u>GCATGCAAGCTAATTCG</u> | for amplifying pPB194 or pDR110 |
| PEB259 | ATGTATACCTCCTTAGTCGACTAAG<u>CTTAATTGTTATCC GCTCACAATTACACACATTATGCCACACCTTGTAGATA</u> | for amplifying pPB194 or pDR110 |
| JLH040 | GCAGGCGAGAAAGGAGAG | ermR cassette forward |
| JLH040a | CGAGGCTCCTGTCACTGC | ermR cassette reverse |
| JLH068 | TTTAGTT**GC**GCCTCAGTTTG<u>AAGCGGGACGGGAATCCG</u> | TlyA$^{K183A}$ catalytic mutant forward |
| JLH068a | CAAACTGAGGC**GC**AACTAAA<u>GCCATGCAGTCGCTGCC</u> | TlyA$^{K183A}$ catalytic mutant reverse |
| JLH069 | AGGTATG**C**GAGCAGG<u>GGCGGCTTAAAGCTCGAAAAAG CGTTG</u> | TlyA$^{V63A}$ SAM binding mutant forward |
| JLH069a | CCTGCTC**G**CATACCT<u>CAGCGGGTTTCCTTTGACAGTTA ACGGAAG</u> | TlyA$^{V63A}$ SAM binding mutant reverse |
| JLH077 | GTGTGTAATTGTGAGCGGATAAC | pPB194 sequencing forward |
| JLH077a | CAAAATCGTCTCCCTCCGTTTG | pPB194 sequencing reverse |
| JLH101 | GGCTTAGC**GC**TCGAAAA<u>AGCGTTGAAGGAATTTCCC</u> | TlyA$^{K69A}$ catalytic mutant forward |
| JLH101A | TTTTCGAGC**GC**TAAGCC<u>GCCCCTGCTCACATAC</u> | TlyA$^{K69A}$ catalytic mutant reverse |
| JLH102 | ACAATTG**C**TGTGTCCTTT<u>ATTTCACTGCGGCTC</u> | TlyA$^{D155A}$ catalytic mutant forward |
| JLH102a | AAAGGACACA**G**CAATTGT<u>GGCAAACTCCGGC</u> | TlyA$^{D155A}$ catalytic mutant reverse |
| JLH103 | CGGAAATATTG**C**GTTTCTCC<u>TTCATTTGCATTGGCCG</u> | TlyA$^{E239A}$ catalytic mutant forward |
| JLH103a | GGAGAAAC**G**CAATATTTCCG<u>TCTCCTCCCGTGATTG</u> | TlyA$^{E239A}$ catalytic mutant reverse |
| JLH105 | CACCG**C**CGGTTTTAC<u>GGACTGCGCTTTGCAAAATGG</u> | TlyA$^{G94A}$ SAM binding mutant forward |
| JLH105a | GTAAAACCG**G**CGGTG<u>GAGGAGCCAATATCAATCATAAT TTTATCTTTGAC</u> | TlyA$^{G94A}$ SAM binding mutant reverse |
| JLH106 | GATATTG**AG**TCCTCCACC<u>GGCGGTTTTACGGACTGCG</u> | TlyA$^{G90E}$ SAM binding mutant forward |
| JLH106a | GGTGGAGGA**CT**CAATATC<u>AATCATAATTTTATCTTTGAC AGAGACGGGAAATTCCTTC</u> | TlyA$^{G90E}$ SAM binding mutant reverse |
| JLH109 | GATATTG**C**CTCCTCCACCGCC<u>G**G**TTTTACGGACTGCG</u> | TlyA$^{G90A+G94A}$ SAM binding mutant forward |
| JLH109a | CGTGGAGGAG**G**CAATATC<u>AATCATAATTTTATCTTTGAC AGAGACGGGAAATTCCTTC</u> | TlyA$^{G90A+G94A}$ SAM binding mutant reverse |
| JLH110 | CGCATGATCTCTTCTTCCG | upstream 817 bp of Phyperspank |
| JLH110a | CAATGGTTCAGATACGACGAC | downstream 752 bp of Phyperspank |

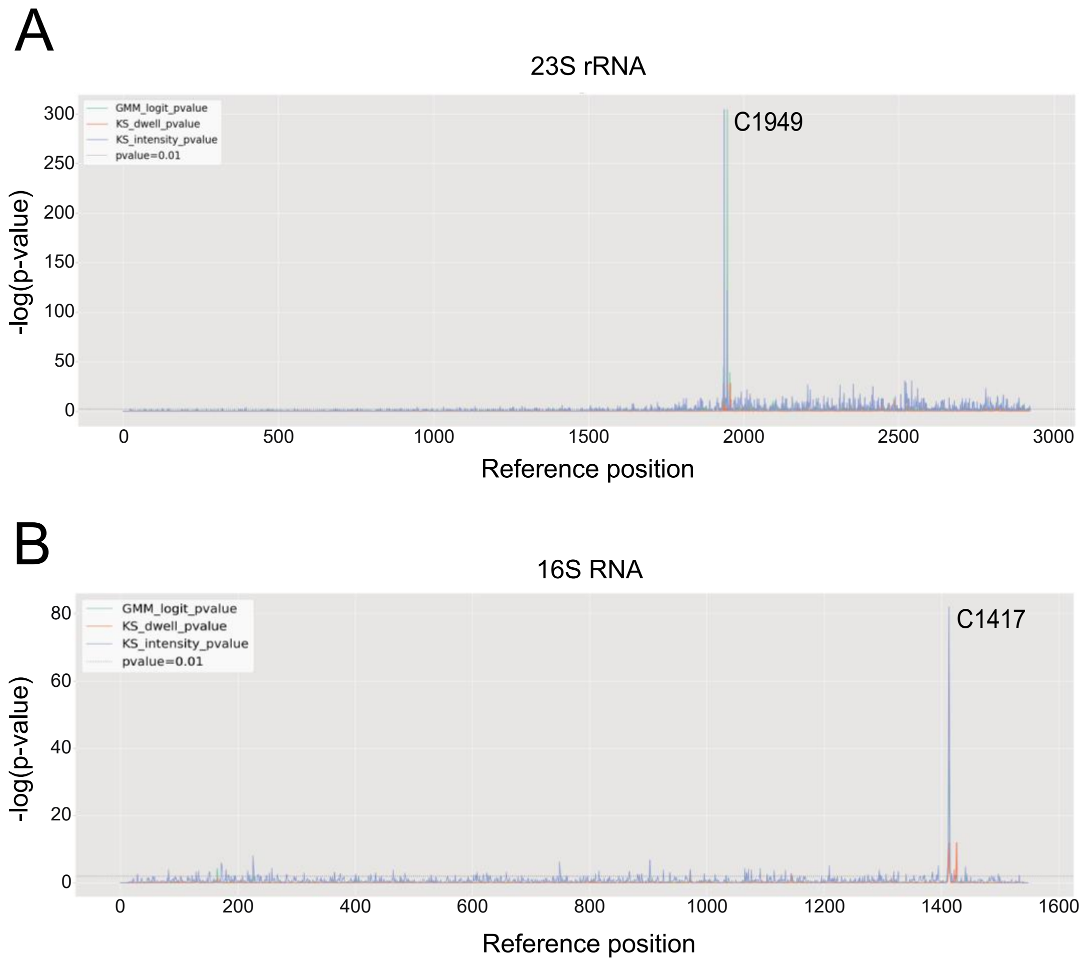


**Figure S1. Nanocompore analysis shows modified cytidine locations in the 23S at C1949 and in the 16S at position C1417.** Shown are p-values from logistic regression log odd ratio (GMM logit) and Kolmolgorov-Smirnov tests for intensity value and dwell time by Nanocompore analysis for every reference position. The p-value peaks identify C1949 for the 23S **(A)** and C1417 for the 16S **(B)**.

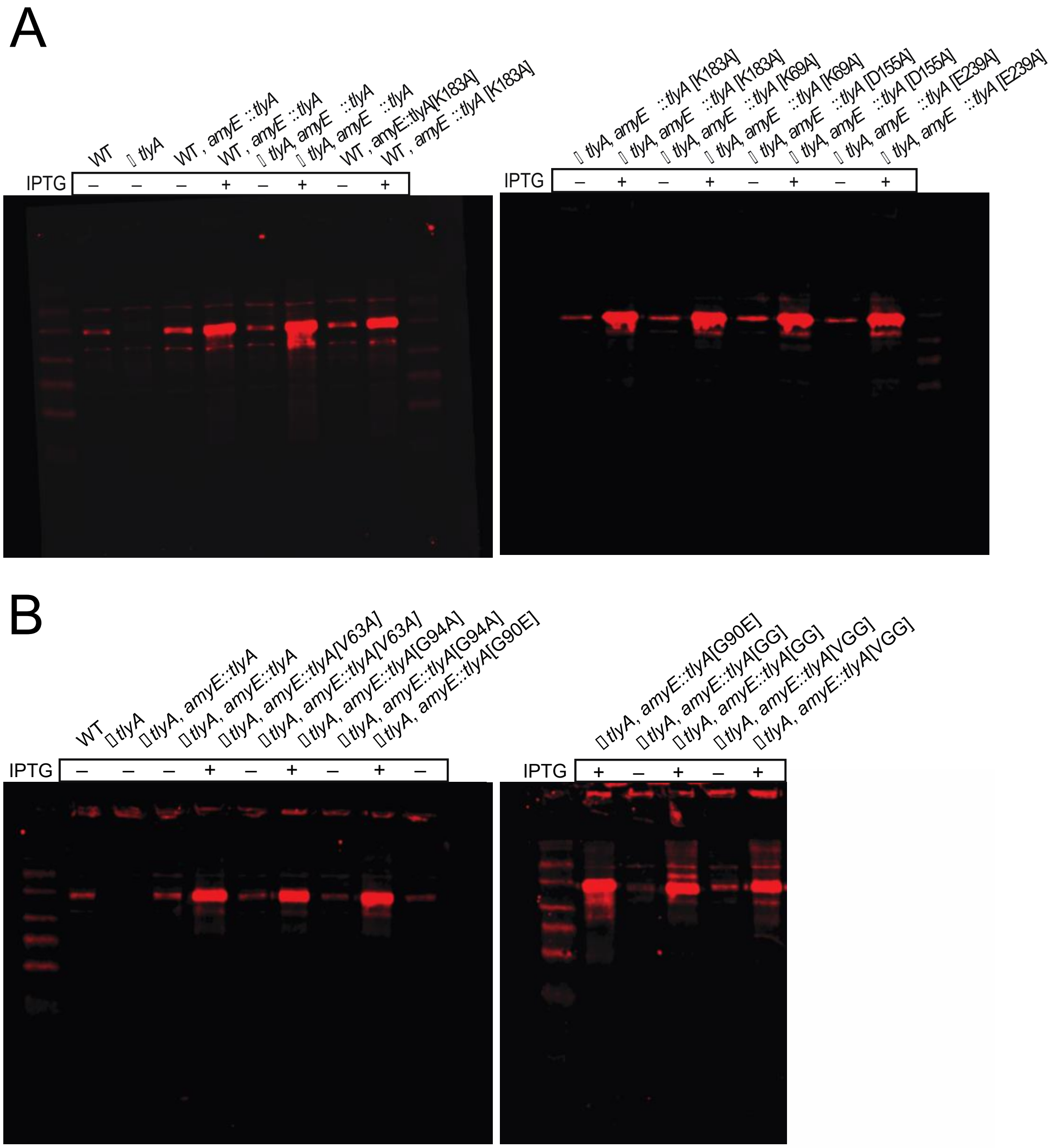


**Figure S2. Full Western blot shows TlyA expression in mutant strains is similar to WT.** Western blot images show mutant TlyA expression and accumula-[illegible] *tlyA* is used as a negative control to show the major immunoreactive band corresponds to TlyA. **(A)** catalytic mutant strains and **(B)** SAM binding mutant strains

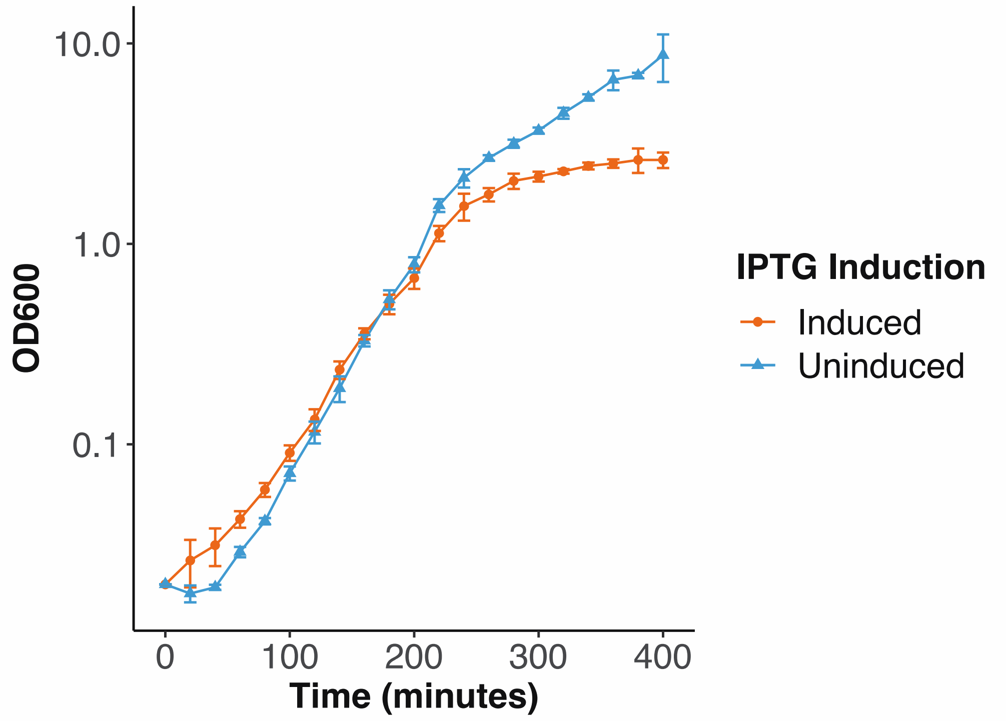


**Figure 3. IPTG induced expression of *tlyA[K183A]* shows reduced growth compared with uninduced expression.** Growth curves show a dominant negative effect when *tlyA*[*K183A*] is induced in a WT background causing a reduction in growth.
